# Trait based assessment of the invasion potential of disease vector mosquitoes

**DOI:** 10.64898/2026.01.05.697723

**Authors:** Rebecca Pabst, Carla A. Sousa, César Capinha

## Abstract

Mosquito-borne diseases pose a growing global health threat, largely driven by the human-mediated spread of vector species beyond their native regions. Although only a few mosquito species historically established populations outside their native ranges, many have expanded rapidly in recent decades. Once established, these invaders are notoriously difficult to control, emphasizing the need for proactive identification before human-mediated spread occurs. Here, we present a framework to anticipate invasion potential for 184 mosquito species of medical importance based on their ecological, life-history, and macroecological traits. We first compiled a comprehensive dataset of 26 traits characterizing each species. We then used random forest models to relate these traits with the probability of species being introduced in new regions (before and after 1950, marking the onset of widespread trade globalization), and of establishment following introduction. Models achieved moderate to good predictive performance (AUC = 0.78-0.85) and revealed that species native to Asia and Australia, adapted to human-made breeding sites, and tolerant of climatic extremes are consistently more likely to be introduced and to establish in non-native regions. Among species with no known invasion history, we identified 24 with higher potential to become future spreaders, of which 17 also exhibit high establishment probabilities (‘high-risk species’). These results show that invasion potential can be inferred, to some extent, from intrinsic species traits and provide a quantitative basis for proactive surveillance, enabling prioritization of species most likely to become introduced in the future.

**Author Summary:** Mosquito-borne diseases threaten more than half of the world’s population and cause over 700,000 deaths each year. Only a small share of mosquito species can spread these diseases, but some of them are moving into new regions where they have never been seen before. The spread of these mosquitoes has led to increasing numbers of locally transmitted outbreaks in regions that previously, or in recent times, had no mosquito-borne diseases. Human activities like trade and travel help mosquitoes spread, and once they arrive, they are extremely difficult to eradicate. Therefore, it is crucial to understand which species may spread in the future and to identify those that should be closely monitored to prevent their introduction and establishment. In this work, we linked species characteristics with their known invasion history to identify the factors driving their introduction and establishment in new regions. We found that species from Asia and Australia, capable of using human-made breeding sites, and tolerant of climatic extremes are most likely to become invaders. We then used these findings to predict which species might spread next. We identified 24 species with high invasion potential, including 17 that also have high chances of establishing once introduced. These results demonstrate that invasion risk can be predicted from measurable species traits, providing a framework to guide early-warning surveillance and prioritize species for monitoring before they begin spreading and become widespread vectors of human disease.

## Introduction

Mosquito-borne diseases are a major threat to global public health, causing more than 700,000 deaths annually and placing over half of the world’s population at risk of infection (1). This risk is largely attributed to a small group of competent vector species capable of transmitting pathogens such as dengue, Zika, chikungunya, malaria, and West Nile virus (2,3). Historically, many of these species maintained relatively restricted distributions. However, in recent decades, the range of several mosquito vectors has expanded at an unprecedented pace. Of the approximately 184 mosquito species, from which human pathogens have been isolated from wild-caught females (4), by now 46 have already been introduced into regions outside their native ranges, with 28 confirmed as having established populations (5). These introductions have expanded the species’ ranges, sometimes into new continents and other fairway regions and enabled local disease transmission in areas considered unsuitable or free of certain diseases, a reminder that while mosquitoes can exist without transmitting pathogens, such pathogens cannot be without mosquitoes (6,7).

Despite receiving increasing attention, the drivers of mosquito introductions and establishment in non-native regions remain poorly understood. Some species, such as *Aedes aegypti*, have expanded globally since the 15th century (8), while others, like *Aedes albopictus*, *Aedes japonicus* and *Anopheles stephensi*, emerged as invasive non-native species (i.e., introduced and established populations outside their native range; cf., 9) only in recent decades (10–12). For many other mosquito species, however, there is no evidence that they were transported by humans or successfully established themselves following introductions. This is intriguing because, in several cases, such species have ecological traits, habitat use, or associations with humans that are broadly similar to those of invasive species (13). Hence, a key question in vector ecology and prevention remains: why are some mosquito species being introduced and becoming established in non-native regions while others remain restricted to their native ranges? Shedding light on drivers of these differences would be of key relevance for supporting surveillance and introduction-prevention efforts, allowing to help identify species more likely to become introduced in the future. Similarly, in a global context where resources for surveillance are limited and the pool of known invasive species is likely incomplete (14), understanding these factors may also help identify introduced or invasive species that remain undetected. To address this question, it is necessary to consider multiple factors shaping the propensity of mosquito species to be introduced and established in non-native regions (15). The disposition to be transported is expected to depend strongly on intrinsic traits that mediate associations with human-traded commodities or transport vectors. Classic examples include oviposition in human-made breeding sites such as used tires and living plant pots, which constitute major pathways for the spread of widespread species such as *Aedes aegypti* and *Ae. albopictus* (16). The breadth of a species’ geographic distribution is also likely to be relevant, with taxa occupying wide native ranges, particularly those overlapping regions of high trade volume and openness, being more frequently exposed to transport opportunities (17). Beyond introduction, establishment success likewise depends on species traits. Water requirements and desiccation resistance influence survival during transit and the prevalence of viable propagules upon arrival (18), while broader environmental tolerances increase the likelihood of encountering suitable conditions for colonization in non-native regions (e.g., salinity 19). Species with higher heat tolerances may also have a competitive advantage under increasingly extreme temperatures (20). In order to provide warning lists to guide preventive biosecurity policies, trait-based approaches have recently emerged as valuable tools to identify species more likely to become introduced and invasive (e.g., 21–23). By integrating variables representing ecological and life-history traits and macroecological patterns into predictive frameworks, these approaches assess which species are more or less likely to enter the global invasion pathway, while also highlighting the traits mediating interspecific variation in invasion potential. For example, Pili et al. (22) developed a global trait-based model and showed that to predict species establishment, both life-history and macroecological traits were important predictors and show that invasion success concentrates in species combining ecological flexibility with high probability of human mediated transport. Similar trait-based assessments for disease vector mosquitoes, would provide critical advances in anticipating future biological invasions and the emergence of vector-borne diseases to protect naïve human populations. Revealing which mosquito species are predisposed to introduction and establishment, would support surveillance planning, refine monitoring priorities and advance our understanding of how ecological specialization, climate tolerance, and human-mediated transport interact to shape global mosquito invasion risk. However, to date, such an assessment has been constrained by the lack of comprehensive trait data, with most compilations restricted to a subset of species of known medical importance (24,25). Here, we take advantage of a newly compiled, extensive database covering 184 mosquito species capable of natural infection with human pathogens to perform a trait-based assessment of species introduction and invasion potential. The trait dataset integrates species ecological, life-history, and macroecological data and to our knowledge is by far, the most taxonomically comprehensive available. Thus, this dataset provides a unique opportunity to test which characteristics of species are associated with a higher or lower invasion potential. Specifically, we combined these data with data from a recent global assessment of mosquitos introduction records and non-native ranges (Pabst et al., 2025) within a machine learning framework to: (i) identify species traits that predict their propensity for introduction and establishment in non-native regions and, (ii) assess which currently unintroduced or non-established species may pose future invasion risks.

## Methods

### 2.1 Trait compilation

Our aim was to estimate the probability of introduction and establishment of mosquito species vectors for human diseases. Species included in the analysis followed Pabst et al. (5) and were limited to mosquito vectors from which human pathogens were isolated from field-caught females, verified by at least two literature sources (4) or by one source plus a disease association recorded in Wilkerson et al. (26). For that purpose, we assembled a database of variables describing ecological and life-history, as well as macroecological traits of these mosquitoes. Traits were selected based on their potential relevance on human-mediated transport of species, their survival during transit, and their establishment in new environments (Table 1). We focused on variables that are relatively stable across environmental gradients and for which information was available for most species, thereby minimizing data gaps and ensuring general applicability. Most ecological and life-history traits (e.g., oviposition behavior, desiccation resistance, salinity tolerance, flight range) were collected through literature research, and each species-trait combination was referenced to a specific literature source (Appendix S1). Macroecological variables (e.g., climatic limits) were derived from species distribution data and spatial environmental layers (CHELSA; 27,28) or obtained from specialized sources like species distribution from Wilkerson et al. (26) and blood meal hosts from Soghigian et al. (29). The rationale for including these derived traits and details on the procedures for their collection are described below.

**Table 1.**
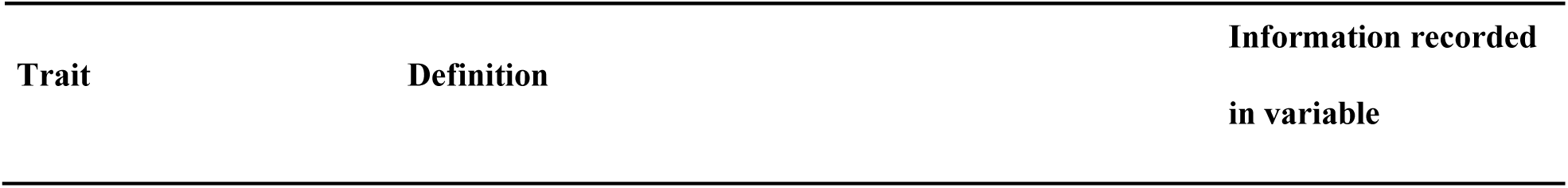
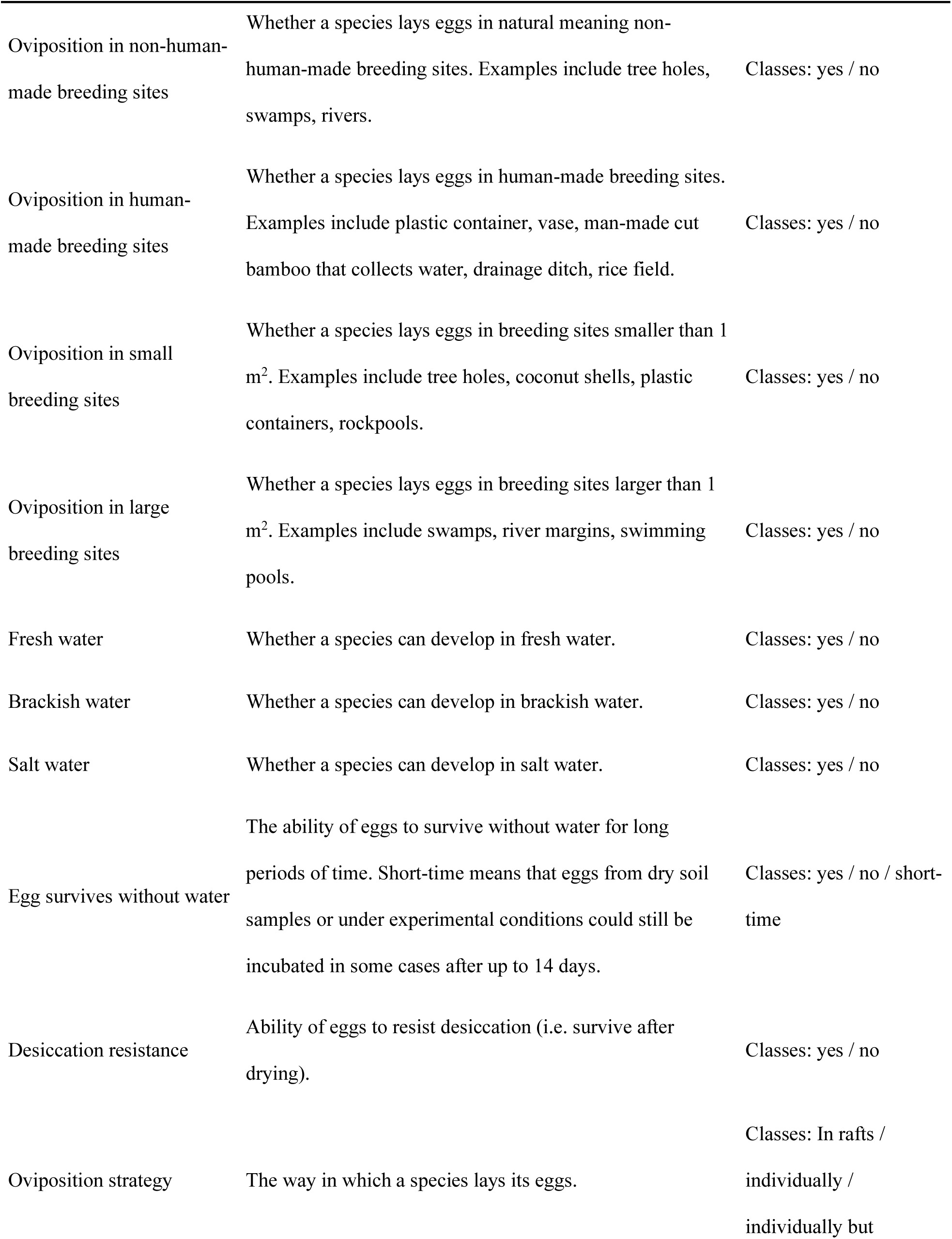
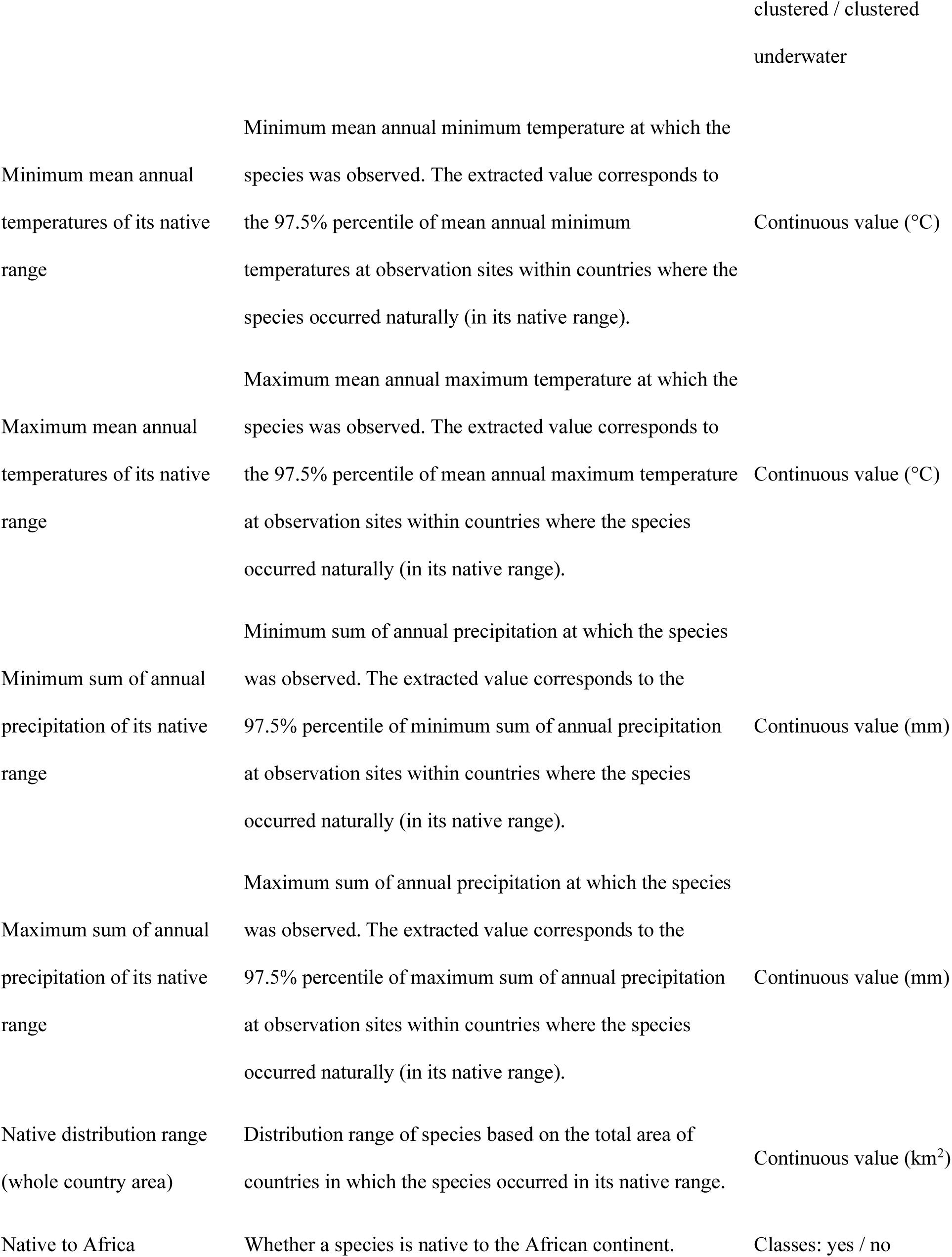
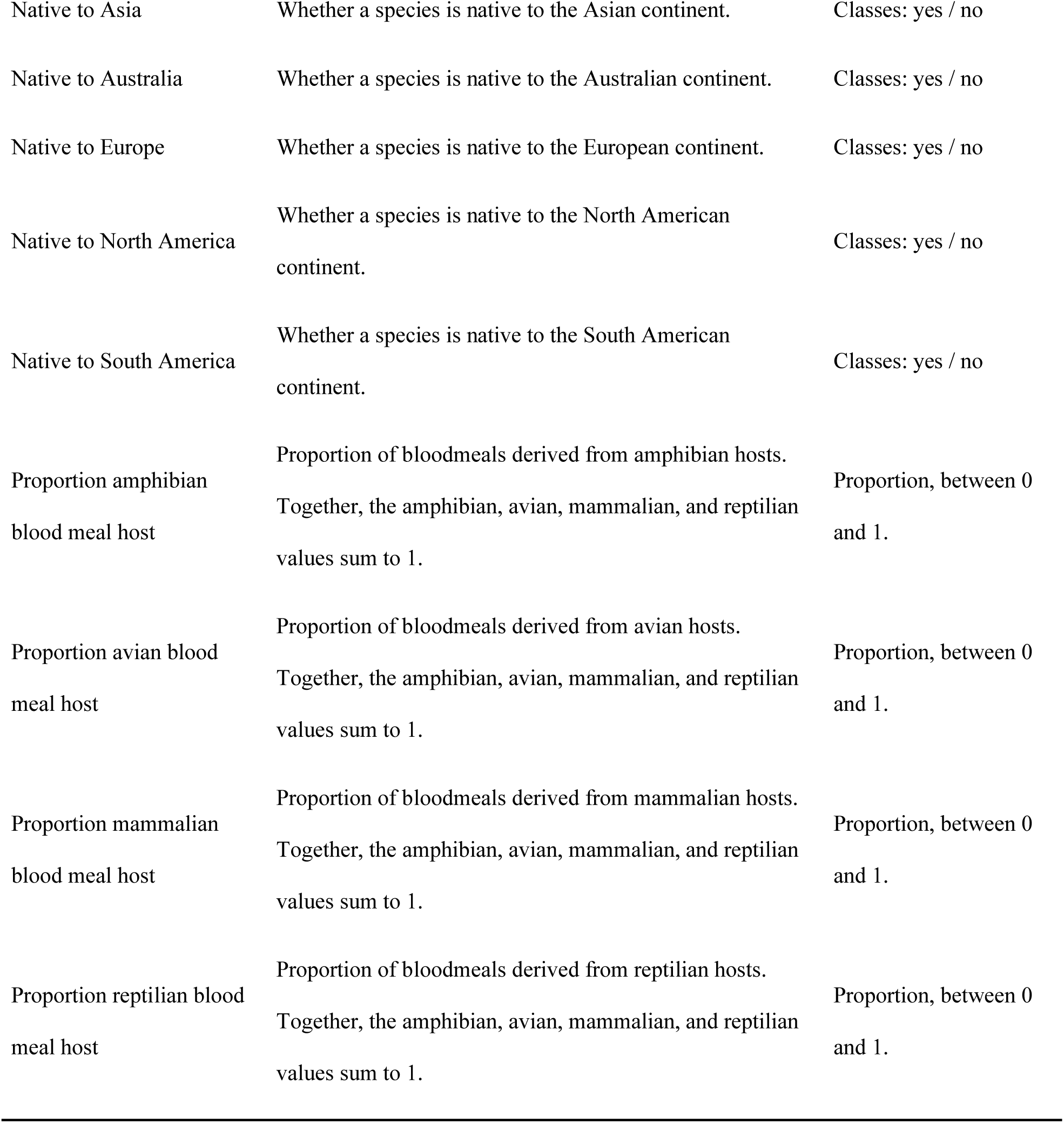
List of species traits collected for the 184 mosquito species of medical importance.

#### 2.1.1 Country wide distribution data and native range delineation

Native ranges were delineated to capture each species’ pre-modern trade-driven distributions and natural ecological context and to avoid temporal overlap between predictors and invasion outcomes. Furthermore, natural distribution determines invasion outcomes due to regional differences in historical trade relations. Country-level data were obtained from Wilkerson et al. (26) and the Walter Reed Biosystematics Unit (30). Each distribution was visually inspected, and only countries within the confirmed native range were retained. Calculated metrics included native-range area (sum of constituent country areas in km²) and native continent(s) as binary variables. Including the latter accounts for the species’ origins.

#### 2.1.2 Climate data

Our trait-based predictive framework requires understanding the climatic tolerances that determine a species’ survival, development, and reproductive cycles. Temperature constrains processes, such as gonotrophic cycle duration, body size, and fecundity (31,32), while precipitation shapes larval habitat availability by influencing standing water body formation and habitat persistence (33). To characterize each species’ climatic niche, we extracted native-range climate values from CHELSA v2.1 layers (27,28) at 30 arc-second resolution (~1 km at the equator). We assembled occurrence records (1980–2024) from GBIF (34) (via the ‘rgbif’ package; Chamberlain et al. 2012) and from VectorMap (30), two global sources of mosquito data. We then cleaned the records for coordinate accuracy, duplicates, and outliers using the ‘CoordinateCleaner’ package (Zizka 2017). Points were restricted to one per grid cell within the species’ native range and overlaid with monthly maximum and minimum temperature, and precipitation layers, to calculate annual averages (temperature) and totals (precipitation). Climatic tolerance thresholds were defined by the 2.5th and 97.5th percentiles to minimize errors from anomalous records caused by errors in georeferencing or species identification.

#### 2.1.3 Host preference

Mosquitoes’ dispersal potential and habitat use are shaped by the availability of their preferred blood meal hosts. Generalist, anthropophilic and mammal-feeding species often occupy urban or peri-urban environments (35), facilitating unintentional human transport, whereas specialised ornithophilic species are typically associated with forested or wetland settings (36). To include this in our model we obtained blood meal host data from Soghigian et al. (29), who compiled molecular blood meal analyses quantifying feeding proportions on mammals, birds, reptiles, and amphibians, providing a standardized host use metric across taxa.

### 2.2 Imputation of missing data

Despite our efforts of data compilation, several species still had incomplete trait information. To ensure the reliability of model estimates, we only included species with at least 75% data completeness and at least 3 unique occurrence points in our modelling (n=169 out of 184). For kept species, the data gaps were imputed using the ‘missForest’ algorithm (37), which iteratively predicts missing values via random forest. To account for imputation uncertainty, we repeated the imputation 100 times, producing 100 complete datasets. Imputation accuracy was evaluated using out-of-bag (OOB) error estimates (normalized root mean square error (NRMSE) for continuous variables; proportion falsely classified (PFC) for categorical variables).

### 2.3 Modeling framework and response variables

We used a random forest (RF) modeling framework to estimate the probability of introduction and establishment of mosquitoes that transmit diseases to humans, based on the fully imputed trait datasets. RF is a nonparametric ensemble learning method that handles continuous, categorical, and partially redundant predictors with high robustness to overfitting (38,39). The algorithm builds multiple decision trees from bootstrap samples and aggregates their predictions, resulting in stable, nonlinear models with strong predictive performance. RF models perform well with sparse or noisy ecological data and offer interpretable measures such as variable importance and partial dependence plots (38,40). These strengths have led to their widespread application in ecological and epidemiological studies (41–44). Our models require two sets of input components: A binary response variable representing a species’ invasion status, and a set of predictor variables describing mosquito ecological, life-history, and macroecological traits. We developed four RF models to assess the invasion potential of mosquito species. The first model estimated the probability of a species being introduced outside its native range, and the second the probability of the species being introduced after 1950. The third and fourth models evaluated the probability of species establishing themselves outside their native range (i.e., forming self-sustaining populations after their introduction), considering either all records, or only those establishments occurring after 1950 respectively. The year 1950 was used as a threshold because it marks the onset of the major acceleration of biological invasions associated with the post-World War II globalization of trade (45). In each model the response variable was binary, coded as “Yes” when species met the criterion and “No” otherwise. Introduction and establishment dates were derived from Pabst et al. (5). Each of these four models was repeated independently 100 times, based on the 100 separate imputation datasets, to generate ensemble predictions that capture prediction uncertainty.

### 2.4 Variable selection

To minimize redundancy and collinearity among predictors, pairwise Pearson correlations were calculated for numerical variables, removing one of each pair with |r| > 0.7 (46). Multicollinearity was further evaluated using the variance inflation factor (VIF) with the ‘vifstep’ function from the ‘*usdm’* R package (47), keeping only variables with a VIF < 5 (48). As a result, we excluded the variables: Oviposition in human-made breeding sites and proportion of avian blood meal hosts. Following this process, in each model we used the same non-redundant combination of continuous, binary, and categorical predictors. The correlation structure of the final numeric variables is shown in Figure 1. In addition to models that included the full set of ecological, life-history, and macroecological traits, we also fitted models using only ecological and life-history variables to assess the stability and consistency of species intrinsic traits as drivers of introduction and establishment. In the main manuscript, we present only the results of the full-variable models, as these performed better and reflect current real-world scenarios; results from the reduced variable set are provided in the Appendix S2.

**Figure 1.**
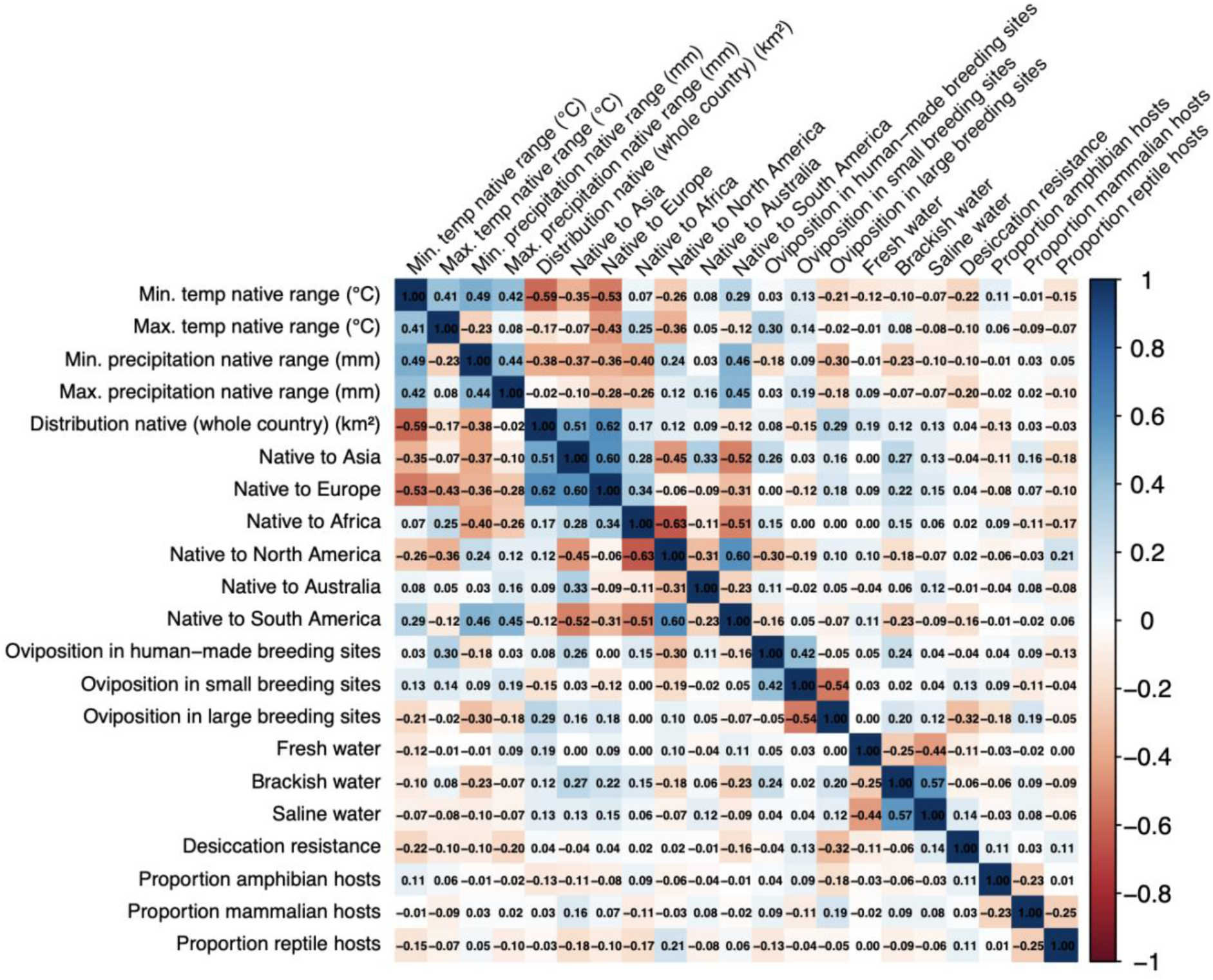
Pairwise Pearson correlation matrix of all numerical predictor variables kept after correlation analysis.

### 2.5 Model fitting and cross-validation

For each response variable, the 100 imputed datasets produced in section 2.2 were used to train 100 separate RF models having the same parameters, implemented in the ‘*ranger’* package (49). Following standard practice, each model was built with 1,000 trees to ensure prediction stability. Each tree used four randomly selected predictors at each split, following the common rule of using the rounded square root of the total number of predictor variables in classification models. A minimum node size of five was specified to reduce overfitting and to ensure that each terminal node contained a sufficient number of observations for stable mean estimates (50).

Model performance was evaluated through leave-one-out cross-validation (LOOCV) (51), in which each species was iteratively withheld from the training set, the model is fitted to the remaining species, and predictions are generated for the excluded species. This was repeated until every species had been left out once, resulting in a full set of out-of-sample predictions. Model performance was evaluated by comparing predicted probabilities with observed outcomes across all species, using the area under the receiver operating characteristic curve (AUC; 52) as a measure of discrimination ability. This approach is robust and mimics a real-world situation where the invasion potential of a species is assessed based on what has been observed for other species. While AUC is a robust, threshold-independent metric for model evaluation and comparison, it does not indicate the probability cutoff at which classification performance is maximized. In our context, identifying such a threshold is important because it allows us to identify which species were correctly or incorrectly classified by the models. To address this, we used the True Skill Statistic (TSS; 53), which provides the probability threshold that maximizes classification accuracy by balancing sensitivity and specificity. The optimal threshold of each RF model was identified by maximizing TSS with the function ‘ecospat.max.tss()’ from the ‘ecospat’ package (54). Predicted probabilities were then averaged across all 100 RF model repetitions to obtain stable ensemble estimates of introduction and establishment probabilities.

### 2.6 Variable importance and trait effects

For each of the four response variables, we additionally ran 100 full RF models with all species to calculate contributions of the predictors using permutation-based importance scores, which represent the decrease in model accuracy after shuffling each variable (38). These values were then averaged to obtain stable estimates of variable importance at the ensemble-level. Variables that showed consistent influence (importance ≥ 0.01 across models) were further explored (50) using partial dependency plots generated from the averaged ensemble predictions, using the FeatureEffect$new() function in the ‘iml’ package (55).

### 2.7 Identifying species with high invasion potential

Using predictions from the LOOCV procedure, we identified species with high potential for future invasion. We first selected taxa not currently introduced but predicted to have a probability of introduction exceeding the TSS-defined threshold. These were considered as potential future spreaders. We then cross-checked these species against predictions from the establishment model and retained those exhibiting simultaneously high probabilities of introduction and establishment, as species of highest invasion concern. All analyses were performed in R (56,57), with scripts and reproducible code provided in the Appendix S3.

## Results

### 3.1 Model performance

A comprehensive dataset was compiled with sufficient data for 169 mosquito species characterized by 24 ecological, life-history and macroecological variables, including eight continuous, three categorical and thirteen one-hot encoded traits. Missing values were imputed with low error, averaging <0.01 % for continuous traits and 3.3 % for categorical traits. Random forest models applied to this dataset yielded cross-validated performances with AUC values indicating moderate (0.78) to good (0.81 and 0.85) performance (Table 2). The introduction models exhibited higher specificity than sensitivity, suggesting stronger performance in identifying non-introduced species, while the establishment models displayed the opposite pattern, reflecting improved recognition of species capable of successful establishment.

**Table 2.** LOOCV random forest model performance metrics (mean ± SD) based on 100 replicates predicting species introduction and establishment probabilities.

| Response | OOB error | Threshold | Accuracy | Sensitivit | Specificity | Precision | F1 | AUC |
| --- | --- | --- | --- | --- | --- | --- | --- | --- |
|  |  | tss |  | y |  |  |  |  |
| Introduced | 0.14±0 | 0.30±0.01 | 0.80±0.01 | 0.78±0.02 | 0.80±0.02 | 0.60±0.02 | 0.65±0.01 | 0.82±0 |
| Introduced after 1950 | 0.15±0 | 0.26±0.03 | 0.76±0.03 | 0.76±0.06 | 0.76±0.05 | 0.51±0.04 | 0.58±0.01 | 0.78±0 |
| Established | 0.11±0 | 0.17±0.01 | 0.77±0.01 | 0.85±0.02 | 0.75±0.02 | 0.42±0.01 | 0.56±0.01 | 0.85±0 |
| Established after 1950 | 0.11±0 | 0.16±0.01 | 0.74±0.01 | 0.84±0.01 | 0.73±0.02 | 0.36±0.01 | 0.50±0.01 | 0.81±0 |

Overall, the establishment models slightly outperformed the introduction models, reflecting greater consistency in identifying traits linked to persistence after arrival. The models correctly identified 35.3 of 46 species known to be introduced beyond their native ranges and 24.1 of 29 established species. For the post-1950 subset, 29.8 of 41 introduced and 21.6 of 26 established species were accurately predicted (Figure 2).

**Figure 2.**
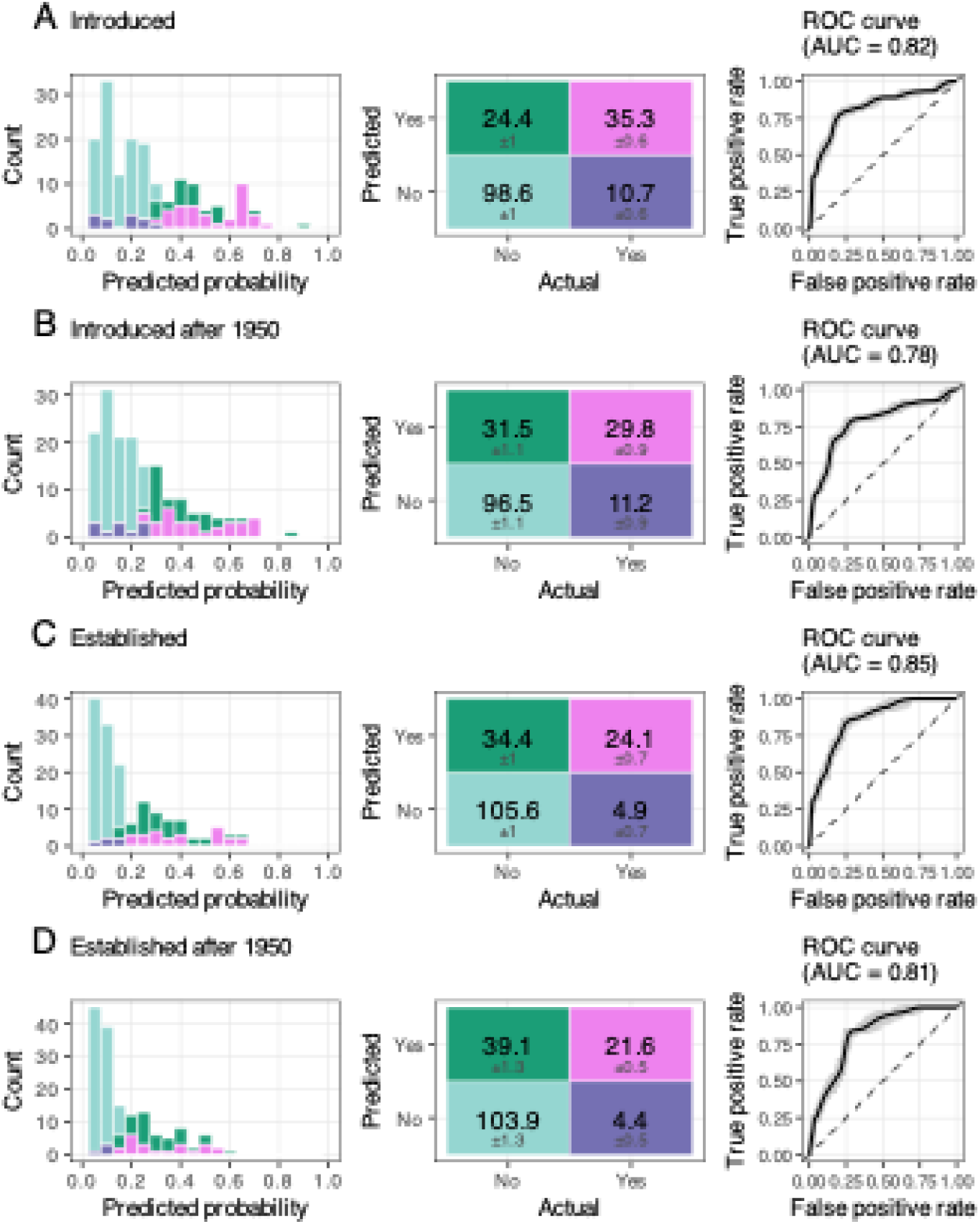
LOOCV random forest model performance. (a) Introduction model (AUC = 0.82); (b) Introduced species after 1950 (AUC = 0.78); (c) Establishment model (AUC = 0.85); (d) Established species after 1950 (AUC = 0.81). Panels show, from left to right, mean predicted probability distributions, mean ± SD confusion matrices, and ROC curves from 100 replicates with the average shown in black.

### 3.2 Variable importance and partial dependency plots

Permutation importance analysis identified native range in Asia as the strongest predictors of mosquito introduction risk, with mean decrease in accuracy of 0.027±0.001 (Figure 3A). Additional influential variables included oviposition in human-made breeding sites (0.018±0.001), native range in Australia (0.016±0.001), native range in North America (0.011±0.001), maximum and minimum precipitation within native ranges (0.015±0.001 and 0.003±0.001), thermal tolerance limits (maximum and minimum temperatures at 0.010±0.001 and 0.011±0.001), and overall native distribution area (0.015±0.001).

**Figure 3.**
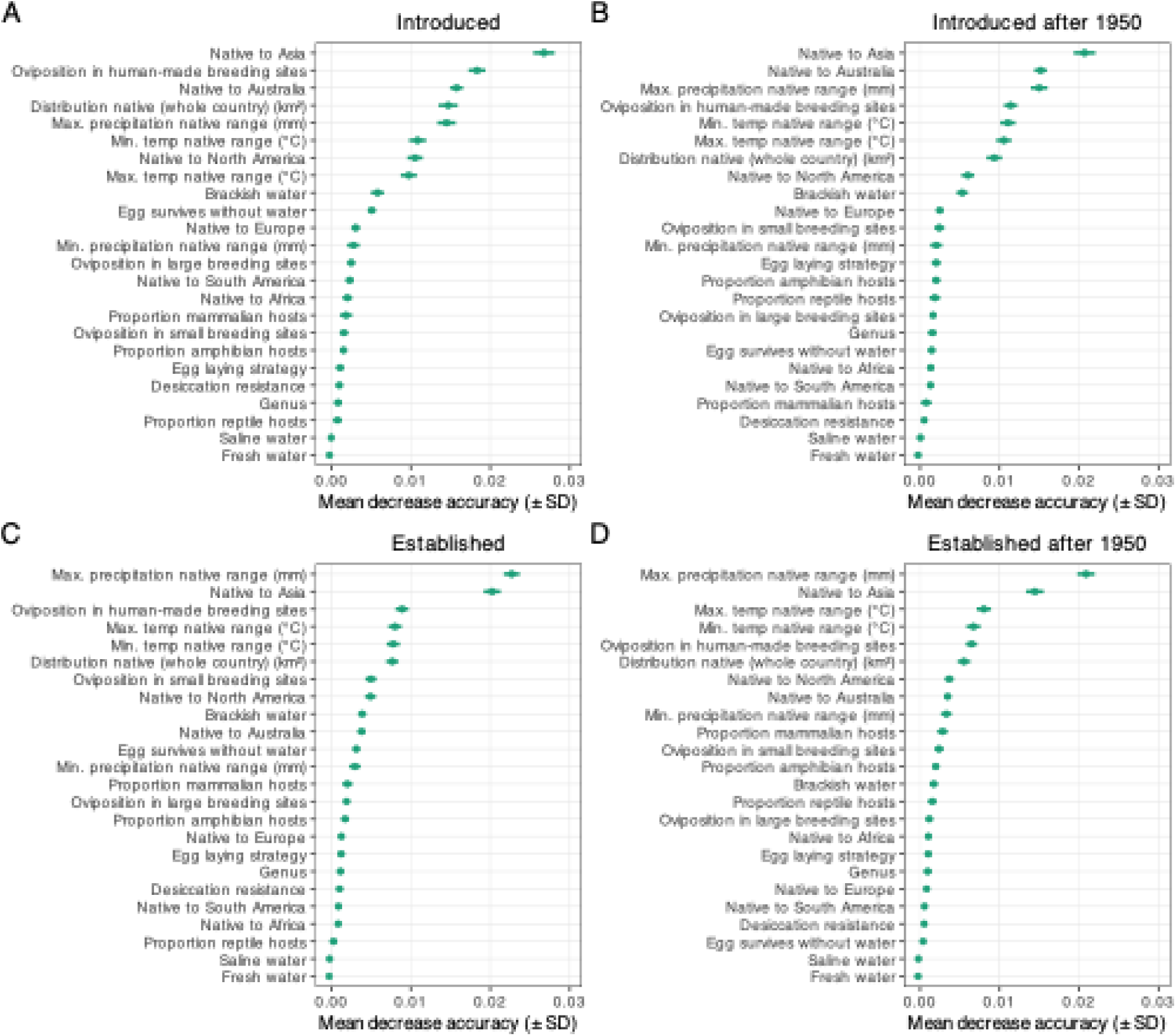
Variable importance rankings from the full random forest models, shown as mean decrease in accuracy (points) ± standard deviation, across 100 model replications. Higher values indicate greater influence in predicting (a) introduction, (b) introduction after 1950, (c) establishment, and (d) establishment after 1950.

In models restricted to species introduced since 1950, origin in Asia remained the predominant predictor (0.021±0.001), along with increased importance of climatic variables such as maximum precipitation, maximum and minimum temperature (0.015±0.001, 0.011±0.001 and 0.011±0.001), consistent with tropical origin of recent introductions, originating in Australia (0.015±0.001) and oviposition in human-made breeding sites (0.011±0.001), (Figure 3B).

For establishment probability in both models, maximum precipitation emerged as the top predictor (0.023±0.001 and 0.021±0.001), followed by Asian origin (0.020±0.001 and 0.014±0.001) (Figures 3C-D). Results based only on ecological and life-history traits were similar in terms of identified and ranking of important variables and the predicted probabilities for the introduction and establishment of species. Major changes were general lower model performance and the emergence of several species predicted as potential spreaders that were not identified by the full models based on all variables (Appendix S2).

Introduction probabilities increased for species native to Asia and Australia, those using human-made breeding sites, those from regions with higher maximum precipitation in their native range and those with wider distribution in original range. The probability decreased for species native to North America. The relationship with minimum temperature was bimodal, with elevated probabilities for species from both cold and warm minimum temperature regimes, whereas species from intermediate temperature ranges were less likely to be introduced (Figure 4A). Although for species introduced after 1950, the maximum temperature of the regions of origin is more important than the distribution area, the patterns remain largely consistent. (Figure 4B).

**Figure 4.**
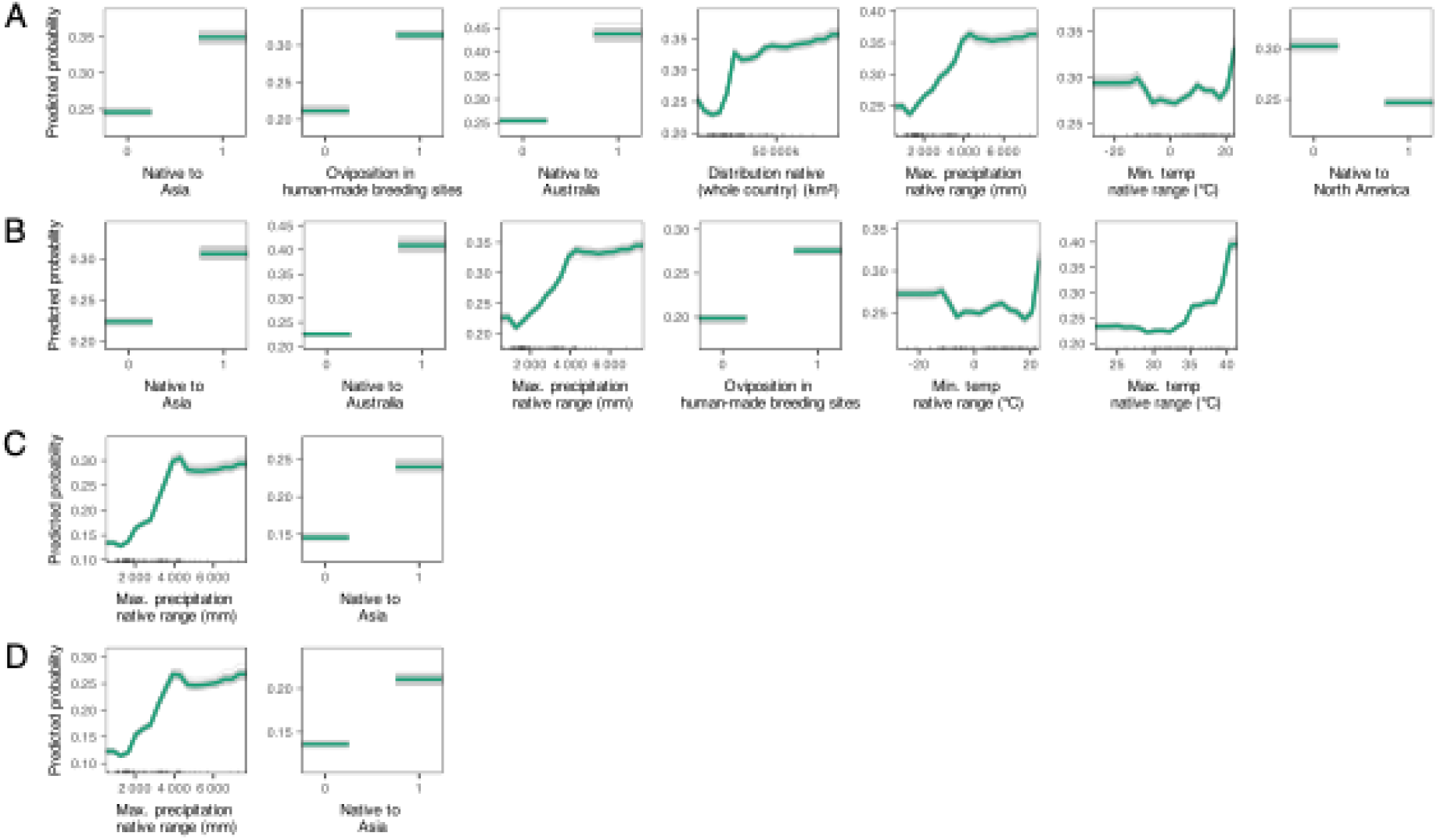
Partial dependence plots illustrating the marginal effects of predictors with importance ≥ 0.01 on predicted probabilities, averaged (green) across 100 random forest model replications (grey). Panels show (a) introduction, (b) introduction after 1950, (c) establishment, and (d) establishment after 1950.

Establishment probability is largely driven by maximum precipitation in their native range and Asian origin (Figures 4C-D).

### 3.3 Predicted spreaders and discrepancies

The introduction model identified 24 mosquito species without known invasion history to have traits consistent with unintentionally transported and introduced species (Table 3). These species share ecological profiles of known invaders, including human altered environments, geographic origin, and climatic characterization of their native range, underscoring their potential as emerging invaders warranting targeted surveillance.

**Table 3.** Predicted potential spreaders. Mosquito species with a model-predicted introduction probability above the average TSS threshold (≥ 0.3) but no confirmed introduction outside their native range. Species in **bold** also show high predicted establishment probabilities, indicating elevated overall invasion risk.

| Species | True Label | Predicted Probability ± SD | Predicted Label |
| --- | --- | --- | --- |
| *Culex vishnui* | No | $0.89 \pm 0.01$ | Yes |
| *Anopheles culicifacies* | No | $0.70 \pm 0.01$ | Yes |
| *Aedes lineatopennis* | No | $0.57 \pm 0.01$ | Yes |
| *Culex univittatus* | No | $0.57 \pm 0.02$ | Yes |
| *Anopheles amictus* | No | $0.56 \pm 0.01$ | Yes |
| *Culex theileri* | No | $0.55 \pm 0.01$ | Yes |
| *Mansonia septempunctata* | No | $0.50 \pm 0.01$ | Yes |
| *Culex torrentium* | No | $0.46 \pm 0.01$ | Yes |
| *Anopheles hyrcanus* | No | $0.45 \pm 0.01$ | Yes |
| *Culex perexiguus* | No | $0.45 \pm 0.02$ | Yes |
| *Culex poicilipes* | No | $0.45 \pm 0.01$ | Yes |
| *Anopheles rufipes* | No | $0.43 \pm 0.02$ | Yes |
| *Anopheles claviger* | No | $0.42 \pm 0.01$ | Yes |
| *Culex nigripalpus* | No | $0.42 \pm 0.02$ | Yes |
| *Anopheles plumbeus* | No | $0.42 \pm 0.01$ | Yes |
| *Culiseta inornata* | No | $0.40 \pm 0.01$ | Yes |
| *Anopheles fluviatilis* | No | $0.40 \pm 0.01$ | Yes |
| *Aedes geniculatus* | No | $0.40 \pm 0.01$ | Yes |
| *Anopheles pseudopunctipennis* | No | $0.37 \pm 0.01$ | Yes |
| *Coquillettidia linealis* | No | $0.36 \pm 0.01$ | Yes |
| *Coquillettidia richiardii* | No | $0.32 \pm 0.01$ | Yes |
| *Anopheles sergentii* | No | $0.32 \pm 0.01$ | Yes |
| *Anopheles pulcherrimus* | No | $0.31 \pm 0.01$ | Yes |
| *Anopheles quadrimaculatus* | No | $0.31 \pm 0.01$ | Yes |

Among them 17 “high-risk” species have attributes consistent with species that also established non-native populations. These include *Culex vishnui, Anopheles culicifacies, Culex univittatus, Aedes lineatopennis, Anopheles amictus, Culex theileri, Mansonia septempunctata, Culex torrentium, Anopheles hyrcanus, Culex perexiguus, Culex poicilipes, Anopheles claviger, Culex nigripalpus, Anopheles plumbeus, Culiseta inornata, Aedes geniculatus,* and *Anopheles pseudopunctipennis*.

On the other hand, seven species that have attributes consistent with accidentally introduced species do not share characteristics with species that were established. Such species may survive transport but fail to overcome ecological or climatic constraints necessary for sustained establishment. These species include *Anopheles rufipes, Anopheles fluviatilis, Coquillettidia linealis, Coquillettidia richiardii, Anopheles sergentii, Anopheles pulcherrimus,* and *Anopheles quadrimaculatus*.

Eleven species with known invasion history were not captured by our introduction model (predicted probabilities < 0.3). The same applies to five species in the establishment model. Most of these species exhibited transient detection or failure to establish self-sustaining populations, suggesting stochastic introduction events or influences from drivers beyond trait-based predictors, such as chance transport or climatic anomalies.

### 3.4 Occurrence of possible spreaders

Below we show the current distributions of species that have been identified as high-risk species obtained from Wilkerson et al. (26), meaning species not yet detected outside their native ranges with high probability of introduction and establishment (Figure 5).

**Figure 5.**
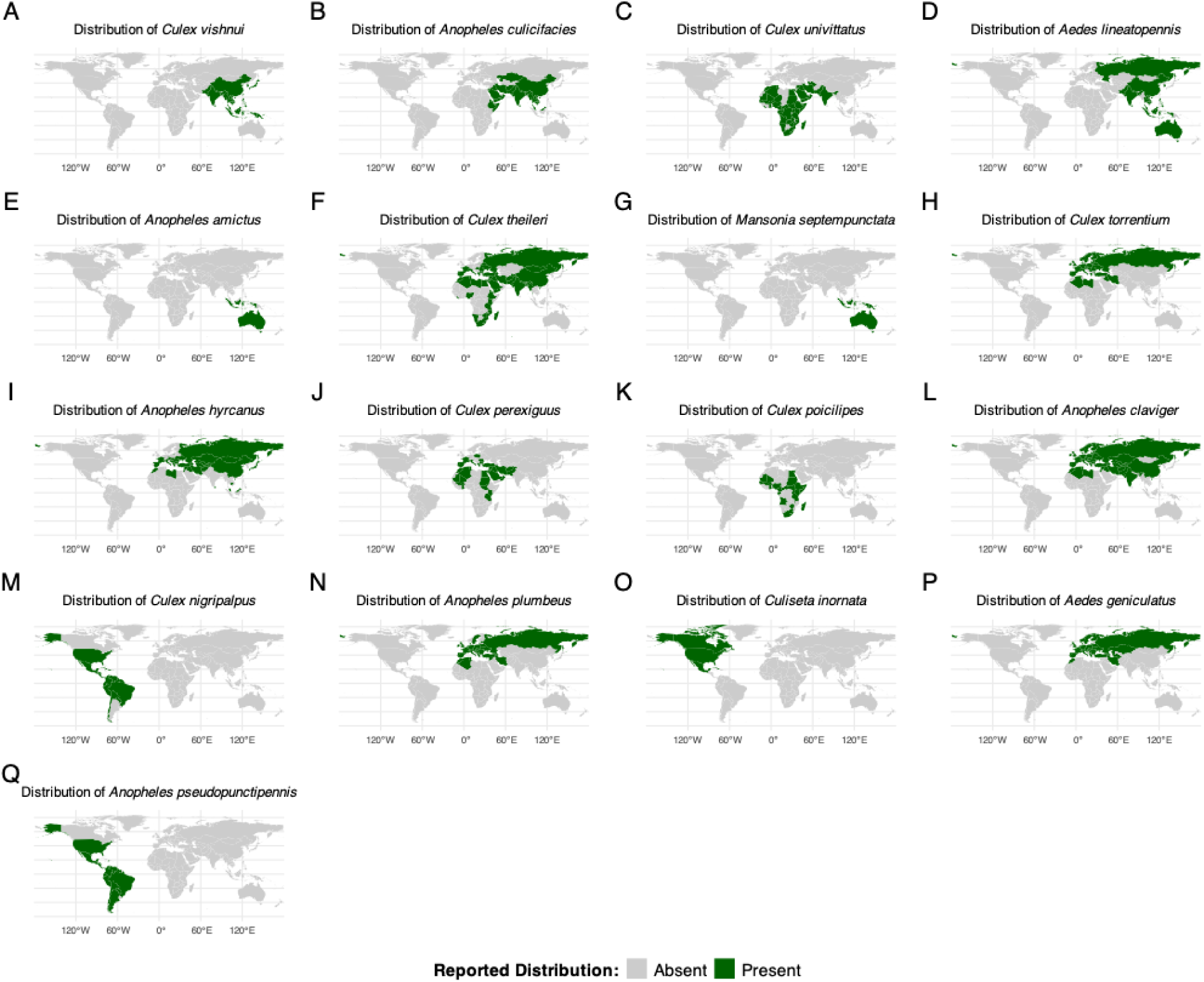
Current distribution of species with high invasion potential, meaning high probability to be introduced as well as to become established. Green shading shows countries where the species are currently reported (26).

## Discussion

Our findings suggest that the introduction and establishment of non-native mosquitoes are driven, to some extent, by some ecological, life-history, and macroecological characteristics. Once established, vector mosquitoes can profoundly alter disease transmission dynamics, or introduce pathogens into previously unaffected regions (3). Responding to their global spread requires predictive tools that go beyond local surveillance and include proactive prevention and risk assessment strategies. Our modelling approach achieved moderate to good predictive ability, stable across model repetitions, which indicates that the invasion potential of species can be inferred to some extent from trait data alone. Unlike previous efforts that modelled the potential distribution of species already introduced (58,59), our framework indicates potential invaders before they spread. Specifically, of the 169 species analyzed, 24 species with no prior invasion history received introduction probabilities equal to or higher than for known introduced species, including 17 species with a high probability of also becoming established. These species are predominantly native to Asia, originated in regions with high precipitation, tend to have broad climatic tolerances, have a wide distribution and are adapted to human-modified environments.

A main result of our analysis is the consistent importance of native biogeographic origin in shaping invasion potential. Species native to Asia and Australia consistently ranked as the most likely to be introduced and to establish outside their native ranges, whereas species from Africa, the Americas or Europe showed substantial lower importance and probabilities. This pattern reflects Asia’s role as principal source region for invasive species overall (60), including invasive insects (61), and recently introduced mosquitoes (5). In Asia, high human population density, intensive containerized trade, and rapid economic expansion create both the propagule pressure and the disturbed habitats that favor human-commensal vectors. Following World War II, used aircraft tires were shipped from Asia and the Pacific region back to the United States as part of postwar recovery and logistics operations because rubber remained a valuable commodity (62), inadvertently facilitating the long-distance transport of mosquito eggs and larvae from that region. Today, the globalization of plant and materials trade continues to provide suitable pathways. For instance, China is by far the leading bamboo exporter (63), and ornamental-plant producers increasingly outsource cultivation to tropical developing countries such as Thailand to reduce labor, land, and infrastructure costs (64). According to the World Bank, China alone accounted for roughly 32% of global container port traffic in 2022 (65). Ongoing large-scale initiatives such as China’s Belt and Road Initiative (BRI) further expand trade and transport networks involving more than 120 countries (66), and the thawing of the Northern Sea Route through the Arctic Ocean (67), potentially opens opportunities for the spread of non-native mosquito species from Asia to Europe. Australia’s significance arises mainly from the fact that most of its native mosquito species were introduced to the Pacific region and New Zealand. Yet, there are several examples of long-distance introductions of mosquitoes native to Australia, including *Aedes vexans* in Hawaii (68), *Anopheles subpictus* in the Netherlands (69), and *Aedes notoscriptus* in California (70). The latter, first detected in Los Angeles County in 2014, shows how integration into global trade networks, combined with climatic similarities between native and introduced regions, can lower environmental barriers and facilitate rapid establishment. In that sense, native continent acts as an integrative factor, capturing evolutionary history, trade exportation intensity, and environmental matching simultaneously. However, global trade networks continuously evolve, potentially reshaping introduction pathways and altering the relative importance of certain origins, routes, and goods (17,45). Nevertheless, our results suggest that the intrinsic traits of a species remain relatively stable predictors, independent of macroecological factors (Appendix S2).

Concerning probability of species introduction, another main predictor identified is the use of human-made breeding sites. Species that oviposit in artificial habitats, plastic vessels, plant pots, rice fields, drainage systems or discarded tires, showed markedly higher probabilities of being transported and introduced than species restricted to natural water bodies. This finding aligns with classic invasion ecology, in which propagule pressure and human association are primary determinants of transport success (15,71). Container breeding has the advantage that eggs and larvae persist in environments closely associated with human activity, trade and travel, from cargo holds to used-tire shipments (72). This pattern, documented e.g., for *Ae. albopictus* and *Ae. japonicus* (10,72,73), is generalizable: nearly all species predicted with high probabilities of both introduction and establishment use human-made breeding sites. This underscores that invasion potential is closely coupled with the degree of adaptation to human-modified landscapes. Global urban expansion and the proliferation of disposable containers continue to multiply these breeding sites, providing abundant breeding opportunities for species with suitable ecological strategies (74,75).

Climatic predictors had a complementary influence, particularly on establishment success. The maximum annual precipitation of the native range was by far the strongest predictor of probability of species establishment, with probabilities rising sharply for species native from areas receiving approximately 1,500 mm yr⁻¹. The relationship between minimum temperature and introduction probability was bimodal: species from both cold and warm native ranges were more likely to be introduced than those from mild climates. This dual pattern might just again stand as a proxy for the species native region, or it suggests that both cold-adapted and tropical mosquitoes possess distinct mechanisms enabling survival during transport, either tolerance to cold or resistance to dehydration. Species from thermally extreme regions on the other hand were more likely to establish, with over half of the successfully established species originating from areas exceeding 34 °C in average maximum temperature. Again, the climate data used here reflect the conditions within the species’ native ranges. However, there are several species were able to establish under environmental conditions that differ from those in their native ranges. This applies, for example, to *Ae. albopictus*, which was initially thought unlikely to establish beyond its native range in tropical and subtropical regions of Southeast Asia, requiring a lengthy adaptation to new ecological conditions and to be constrained by competition with local mosquito species (76). Nevertheless, it rapidly adapted to colder conditions (77,78), outperformed presumed competitors (79,80) and expanded into temperate regions worldwide. Hence, predictions of invasion and establishment potential based solely on native-range climates should be interpreted with caution.

Finally, we also found that widely distributed species are also more likely to be introduced into new regions. This may translate into increased propagule pressure as greater distribution increases the chances of introduction. This was however not observed for established species, which could indicate a filter for species establishment that is based more on intrinsic traits or adaptation to new environments (20,81).

Overall, our results indicate that mosquito invasion potential is best explained by the interaction between life-history flexibility and macroecological breadth. Adaptation to human environments and occurrence in regions that are export hubs for commodities associated with human-made oviposition sites that promote dispersal and introduction. In turn, broad climatic tolerance, particularly to extremes of rainfall and temperature, determines whether colonization results in successful establishment. Together, these traits define a functional profile of likely invaders: human-adapted species, often native to Asia and/or Australia, capable of exploiting human-made breeding sites, tolerant of precipitation and thermal extremes and with a wide distribution. These findings extend previous qualitative insights (18) to a quantitative global framework, go beyond the few medically very important vectors (e.g., 59,72,82,83), and provide insights into the invasive potential of lesser-known species. The results also demonstrate that invasion potential in mosquitoes can be forecasted to some extent from measurable ecological, life-history, and macroecological traits.

Beyond identifying traits of importance, our models identified 24 species with no recorded invasion history but trait profiles closely matching those of known invaders. Of these, 17 species were also predicted with high probabilities of establishment, marking them as priority candidates for early surveillance. For example, *Culex vishnui,* the species with the highest predicted introduction probability and likely to establish is native to South and East Asia, where it thrives in rice fields, ground pools, and small artificial containers (26). It has been found naturally infected with multiple arboviruses, including Japanese encephalitis and West Nile virus (84,85).

*Anopheles culicifacies*, ranking second in our high risk species list, is an important malaria vector, particularly in South and Southwest Asia. (86). Predominantly anthropophilic but occasionally zoophilic, it can take multiple blood meals per gonotrophic cycle (26) increasing its potential for pathogen transmission.

*Aedes lineatopennis*, native to the Australasian and Oriental regions, was first described from the Philippines (26,87). It has been found naturally infected with Japanese encephalitis (88) and Middelburg virus (89), and is considered a potential vector for Ross River and Murray Valley encephalitis viruses (90,91).

Our results provide valuable insights, but their interpretation should also take certain considerations into account. First, trait data for some mosquitoes remains partly incomplete. Similar to other studies (21,22,92), we used a trait imputation approach to overcome this limitation. To minimize bias, we included only species with at least 75% trait completeness, and only variables that were known for at least 75% of the species. Nonetheless, the imputation process may obscure subtle ecological differences that influence invasion potential. Second, because our approach is species trait–centric, it does not account for the local conditions species encounter at introduction sites, such as climate, biotic interactions, or vector control measures that can influence invasion outcomes (93,94). Furthermore, environmental change can modify the current distribution of suitable habitats, impairing species establishment success. Thus, our framework should be regarded as an initial profiling tool for species with invasion potential, to be complemented by geographically explicit assessments of establishment risk. Finally, it is important to recognize that the predictors used in our models are not static. Global trade networks may continue to evolve and other regions of origin may become sources for species exports. However, results based solely on ecological and life-history traits gave similar predicted probabilities of species introduction and establishment showing the resilience of our findings (Appendix S2). Thus, although our results may be temporally bounded, our framework provides a robust baseline for identifying species with elevated invasion potential under present-day ecological and trade conditions and are most relevant for near-term invasion assessments.

## 5 Conclusion

Overall, our results indicate that the invasion potential of mosquitoes can be partially predicted from intrinsic biological and ecological traits alone. Species displaying traits like being native to Asia and Australia, using human-made breeding sites, from regions with climate extremes can be expected to be those posing higher risk of invasion in non-native regions in the near future. By identifying these species and their characteristic profiles, our approach underscores the potential for proactive, trait-based surveillance strategies that extend beyond the currently recognized vector species. Trait-based frameworks such as the one presented here can support early-warning systems, guide allocation of surveillance resources, and ultimately reduce the risk of novel vector-borne disease emergence. Once invasive mosquitoes succeed in establishing populations, their control and eradication become exceedingly challenging (95,96). In this context, preventing introductions remains the most efficient and economically viable strategy.

## Supporting information

Appendix S2: Additional model results.

## Acknowledgments

RP and CAS gratefully acknowledge the support of the Portuguese Foundation for Science and Technology (FCT) for funds to the R&D Unit Global Health and Tropical Medicine (UIDB/04413/2025) and the Associated Laboratory in Translation and Innovation Towards Global Health REAL (LA/P/0117/2020). RP acknowledges funding from FCT (PRT/BD/153694/2021; https://doi.org/10.54499/PRT/BD/153694/2021) and thanks the AIR Center for their support. CC acknowledges funding from FCT through InvaSTOP grant (https://doi.org/10.54499/2023.12533.PEX) and the support from FCT through funds to CEG/IGOT Research Unit (https://doi.org/10.54499/UID/00295/2025).

## Supporting information captions

**Appendix S1: Traits data set.** Data set generated with all species traits combinations and corresponding sources of information will be made available upon acceptance of article

**Appendix S2: Additional model results.** Model results of models using only ecological and life-history traits, not macroecological traits, 3 Figures, 2 Tables

**Appendix S3: R Code for analysis.** Compressed folder containing the R script used to perform the analyses, along with a simplified version of the input dataset will be made available upon acceptance of the article. The original global mosquito introduction dataset can be accessed at: https://doi.org/10.5281/zenodo.15731141.

