## Appendix S2: Additional model results. for "Trait based assessment of the invasion potential of disease vector mosquitoes"

### Appendix S2 for:

### Trait based assessment of the invasion potential of disease vector mosquitoes

**Authors**

Rebecca Pabst[1*], Carla A. Sousa[1], César Capinha[2,3]

**Affiliations**

[1] Global Health and Tropical Medicine, GHTM, LA-REAL, Institute of Hygiene and Tropical Medicine, IHMT, NOVA University Lisbon, Lisbon, Portugal.

[2] Centre of Geographical Studies, Institute of Geography and Spatial Planning, University of Lisbon, Lisboa, Portugal.

[3] Associate Laboratory TERRA, Lisboa, Portugal.

3 Figures

2 Tables

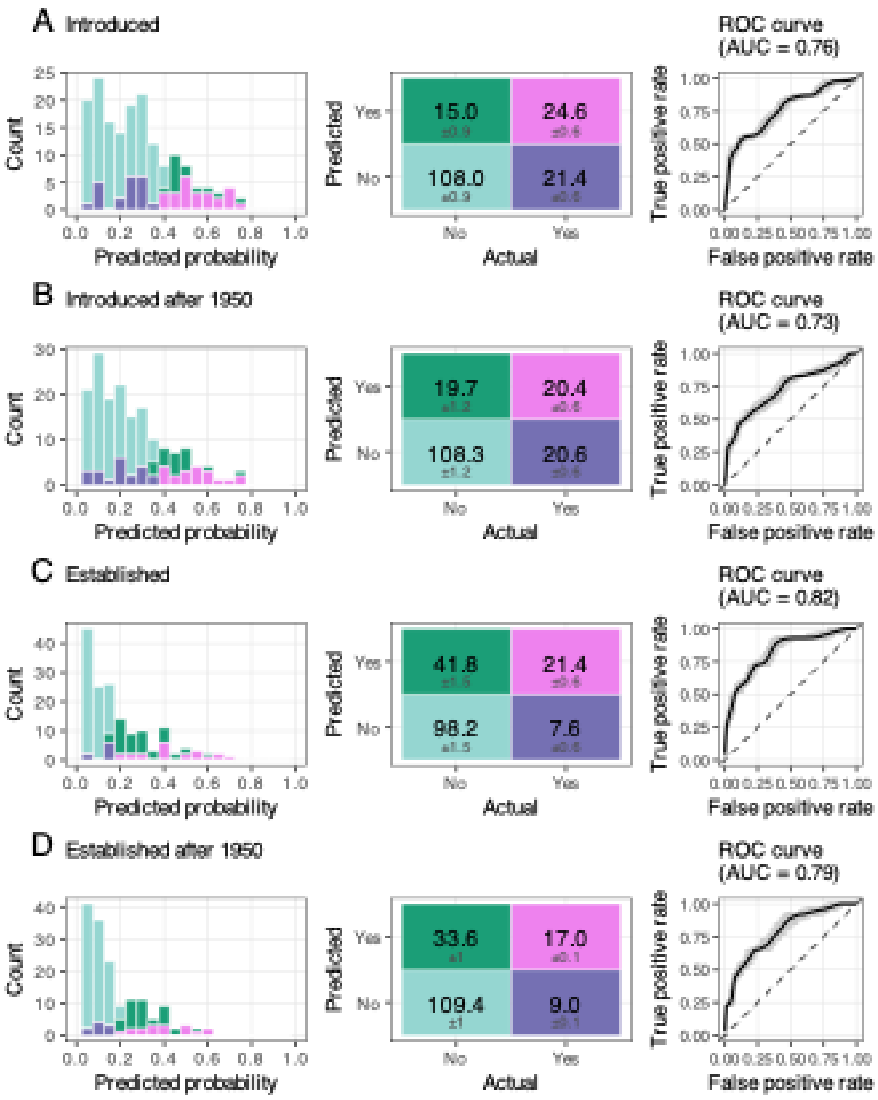


**Appendix S2 Figure 1:** Model performance of LOOCV random forest models including only ecological and life-history traits. (a) Introduction model (AUC = 0.76); (b) Introduced species after 1950 (AUC = 0.73); (c) Establishment model (AUC = 0.82); (d) Established species after 1950 (AUC = 0.79). Panels show, from left to right, mean predicted probability distributions, mean ± SD confusion matrices, and ROC curves from 100 replicates with the average shown in black.

**Appendix S2 Table 1:** LOOCV random forest model performance metrics (mean ± SD) based on 100 replicates predicting species introduction and establishment probabilities, including only ecological and life-history traits as predictors.

| Response | OOB error | Threshold tss | Accuracy | Sensitivity | Specificity | Precision | F1 | AUC |
|---|---|---|---|---|---|---|---|---|
| Introduced | 0.16±0 | 0.39±0.02 | 0.79±0.02 | 0.54±0.03 | 0.88±0.04 | 0.63±0.04 | 0.58±0.01 | 0.76±0 |
| Introduced after 1950 | 0.16±0 | 0.35±0.05 | 0.77±0.04 | 0.54±0.08 | 0.84±0.08 | 0.54±0.07 | 0.53±0.01 | 0.73±0 |
| Established | 0.11±0 | 0.17±0.05 | 0.71±0.06 | 0.83±0.10 | 0.69±0.09 | 0.37±0.05 | 0.50±0.03 | 0.82±0 |
| Established after 1950 | 0.11±0 | 0.20±0.05 | 0.74±0.07 | 0.70±0.10 | 0.75±0.10 | 0.35±0.05 | 0.46±0.03 | 0.79±0 |

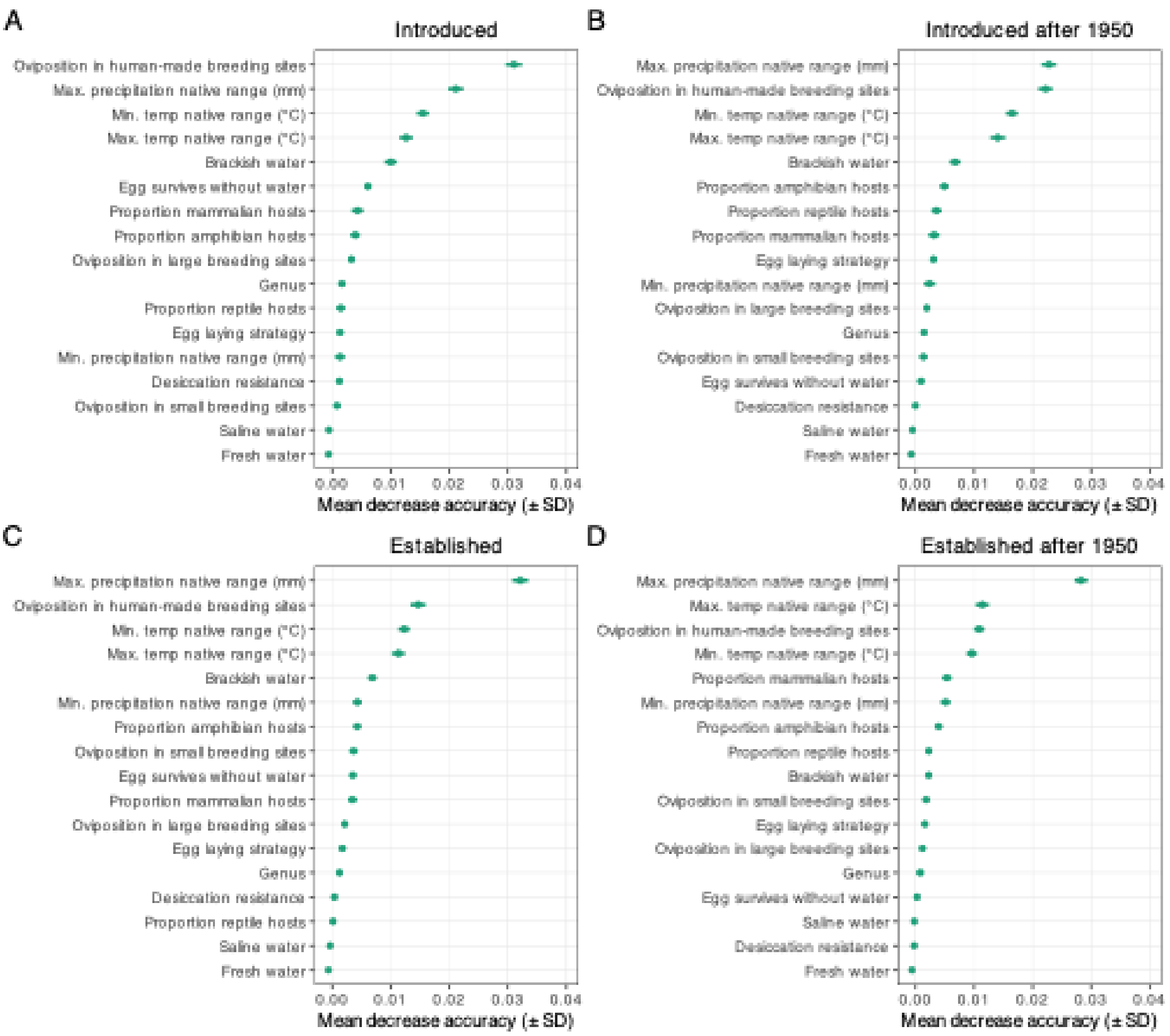


**Appendix S2 Figure 2:** Variable importance rankings from the random forest models including only ecological and life-history traits, shown as mean decrease in accuracy (points) ± standard deviation, across 100 model replications. Higher values indicate greater influence in predicting (a) introduction, (b) introduction after 1950, (c) establishment, and (d) establishment after 1950.

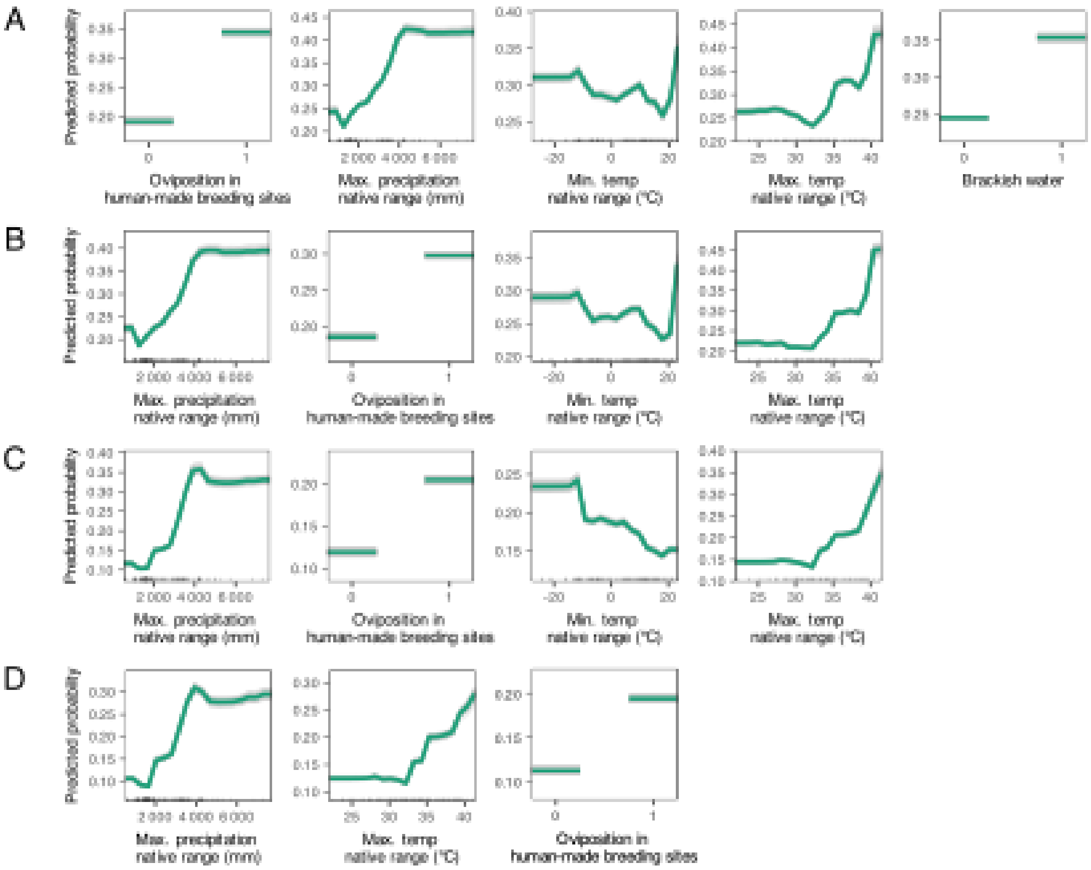


**Appendix S2 Figure 3:** Partial dependence plots of models fitted only with ecological and life-history traits, illustrating the marginal effects of predictors with importance ≥ 0.01 on predicted probabilities, averaged (green) across 100 random forest model replications (grey). Panels show (a) introduction, (b) introduction after 1950, (c) establishment, and (d) establishment after 1950.

**Appendix S2 Table 2:** Predicted potential spreaders based only on ecological and life-history traits. Listed are mosquito species with an introduction probability, as well as establishment probability above the average TSS threshold, but with no confirmed introductions outside their native range. Species shown in **bold** were not identified as high-risk in the full model that also included macroecological variables.

| Species | True Label | Predicted Probability ± SD | Predicted Label |
|---|---|---|---|
| *Culex vishnui* | No | 0.76 ± 0.01 | Yes |
| *Culex nigripalpus* | No | 0.63 ± 0.01 | Yes |
| *Anopheles culicifacies* | No | 0.61 ± 0.01 | Yes |
| ***Anopheles nuneztovari*** | No | 0.57 ± 0.02 | Yes |

| | | | |
|---|---|---|---|
| *Anopheles pseudopunctipennis* | No | 0.48 ± 0.01 | Yes |
| ***Anopheles grabhamii*** | No | 0.48 ± 0.02 | Yes |
| ***Aedes africanus*** | No | 0.47 ± 0.02 | Yes |
| *Culex perexiguus* | No | 0.47 ± 0.02 | Yes |
| *Culex theileri* | No | 0.46 ± 0.01 | Yes |
| *Culiseta inornata* | No | 0.46 ± 0.01 | Yes |
| *Culex torrentium* | No | 0.46 ± 0.02 | Yes |
| ***Aedes taeniorhynchus*** | No | 0.45 ± 0.02 | Yes |
| *Culex univittatus* | No | 0.44 ± 0.02 | Yes |
| *Aedes geniculatus* | No | 0.40 ± 0.02 | Yes |
